# mRNA based SARS-CoV-2 vaccine candidate CVnCoV induces high levels of virus neutralizing antibodies and mediates protection in rodents

**DOI:** 10.1101/2020.10.23.351775

**Authors:** Susanne Rauch, Nicole Roth, Kim Schwendt, Mariola Fotin-Mleczek, Stefan O. Mueller, Benjamin Petsch

## Abstract

mRNA technologies have recently proven clinical efficacy against coronavirus disease 2019 (COVID-19) and are among the most promising technologies to address the current pandemic. Here, we show preclinical data for our clinical candidate CVnCoV, a lipid nanoparticle encapsulated mRNA vaccine that encodes full length, pre-fusion stabilised severe acute respiratory syndrome coronavirus-2 (SARS-CoV-2) Spike protein. In contrast to previously published approaches, CVnCoV is exclusively composed of naturally occurring nucleotides. Immunisation with CVnCoV induced strong humoral responses with high titres of virus neutralizing antibodies and robust T cell responses. CVnCoV vaccination protected hamsters from challenge with wild type SARS-CoV-2, demonstrated by the absence of viral replication in the lungs. Hamsters vaccinated with a suboptimal dose of CVnCoV leading to breakthrough viral replication exhibited no evidence of vaccine enhanced disease. Overall, data presented here provide evidence that CVnCoV represents a potent and safe vaccine candidate against SARS-CoV-2.

## Introduction

The global COVID-19 pandemic has highlighted the need for novel technologies that allow rapid development and production of human vaccines against newly emerging infectious pathogens. Following pioneering work using mRNA formulated with protamine to target tumours^1, 2, 3, 4^, CureVac has established that mRNA elicits immune responses against target antigens as a prophylactic vaccine^5,6,7,8,9^. CureVac’s proprietary mRNA technology is designed to rapidly identify, produce, and test stable and immunogenic mRNA molecules^10^. Following preclinical proof of concept with rabies glycoprotein RABV-G mRNA, formulated with protamine^8,7^, a first-in-human study showed that immune responses are elicited in adult volunteers, although protective titres could only be induced when specialised injection devices were used^9^. Further research has demonstrated that RABV-G mRNA encapsulated in lipid nanoparticles (LNP) overcomes these deficiencies and significantly improves vaccine efficacy in animal models^6^, and in human volunteers^11^.

mRNA technology is now the basis for several severe acute respiratory syndrome coronavirus 2 (SARS-CoV-2) vaccine candidates^12; 13,14,15,16^. The main antigenic target of SARS-CoV-2 is the glycosylated spike protein (S) which interacts with human angiotensin-converting enzyme 2 (ACE2). Consistent with the mode of action of SARS-CoV, which first emerged in 2002-2003^17^, ACE2 binding allows cellular entry of the virus^18,19,20^. S is a trimeric glycoprotein complex located on the viral surface and is a critical target for viral neutralizing antibodies^21^. Each monomer consists of two domains, S1 and S2 that act separately to mediate viral binding and fusion to the host cell membrane, respectively. The S1 domain interacts with cell-surface receptors through a receptor binding domain (RBD) and monoclonal antibodies (mAb) against the RBD possess neutralizing capacity^22^. Fusion with the membrane through S1 leads to a conformational change in the spike protein, proteolytic cleavage of the S1 and S2 domains, and ultimately, viral uptake and replication^21^,^23^.

CureVac has applied its mRNA technology to the rapid development of CVnCoV, a SARS-CoV-2 vaccine designed for maximal protein expression and balanced immune activation. CVnCoV is comprised of LNP-formulated, non-chemically modified, sequence engineered mRNA encoding full length S protein with two proline mutations (S-2P). These mutations stabilize protein conformation as previously reported for Middle East respiratory syndrome coronavirus (MERS-CoV)^24^ and SARS-CoV^25^. Here we describe the immunogenicity and protective efficacy of CVnCoV in preclinical studies in rodents. Protective efficacy was assessed in Syrian hamsters, one of the recognized and accepted models to investigate human-relevant immunogenicity and pathogenesis^26^. Hamsters are susceptible to wild-type SARS-CoV-2 infection, resulting in high levels of virus replication and histopathological changes in viral target organs comparable to mild to moderate human lung disease pathology. Studies shown here enabled the start of CVnCoV clinical development^27^, currently in phase 2b/3 clinical studies.

## Results

CVnCoV encodes for full-length SARS-CoV-2 S protein with intact S1/S2 cleavage site and transmembrane domain, as well as K_986_P and V_987_P mutations^24,25^ (S-2P) (Figure 1A). Nonencapsulated SARS-CoV-2 S-2P mRNA translated in a cell free *in vitro* system yielded a 140 kDa protein, representing uncleaved full length S-2P (Figure 1B). Efficient cleavage of the S-2P protein in cell culture was demonstrated by Western blot analysis of mRNA transfected cells, using an antibody directed against the S2 portion of the protein^28, 20^. This analysis showed the generation of two main bands of approx. 90 kDa and 180 kDa, likely reflecting glycosylated forms of unprocessed S protein (S0) and the cleaved S2 subunit (Figure 1C). In agreement with an intact signal peptide and transmembrane domain, further flow cytometric analyses proved that a large proportion of mRNA encoded SARS-CoV-2 S localised to the surface of mRNA transfected cells.

**Figure 1:**
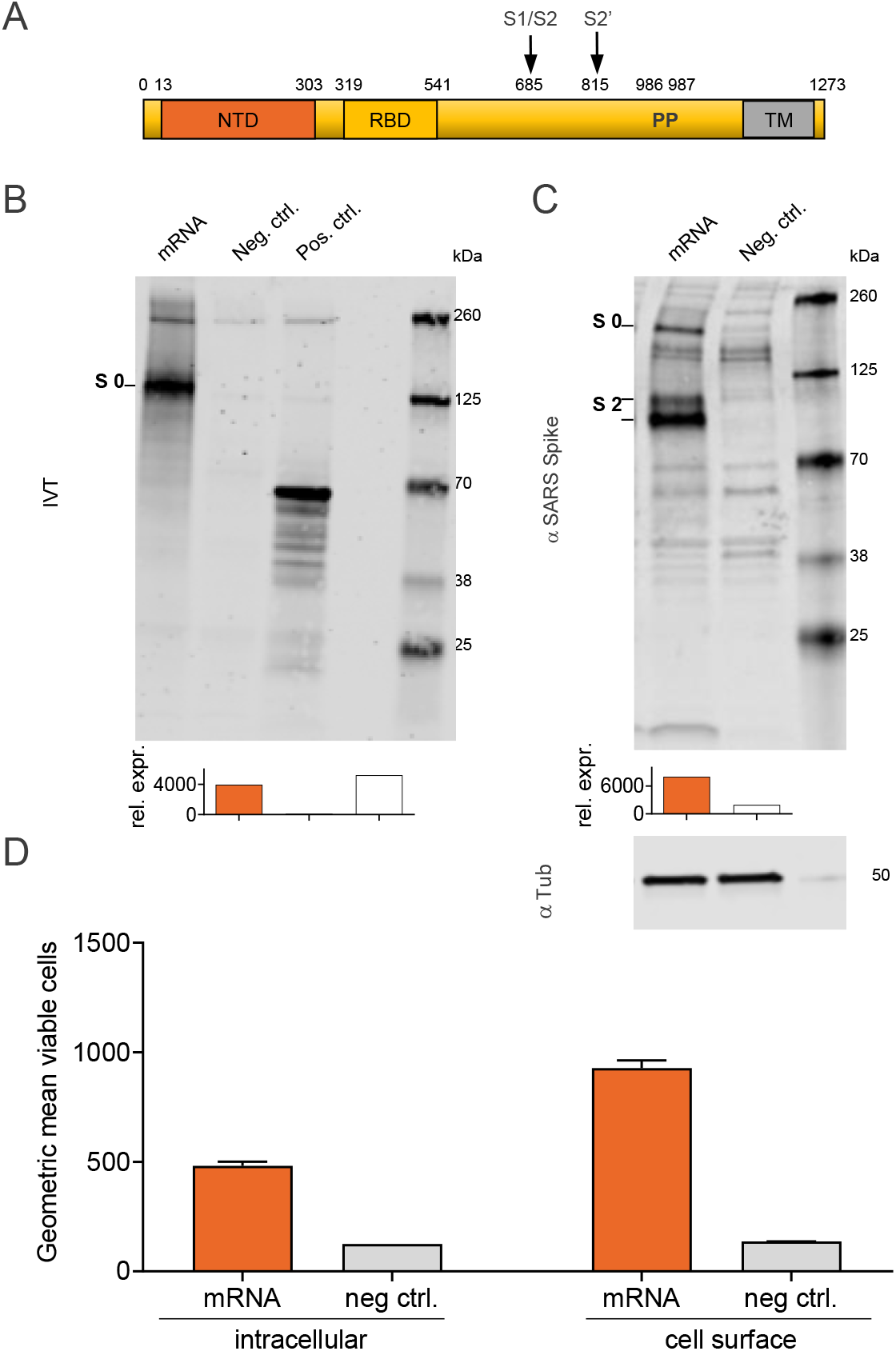
Protein translated from CVnCoV is cleaved, post-translationally modified and presented on the cell surface. (A) Schematic drawing of SARS-CoV-2 S-2P encoded by CVnCoV (B) *In vitro* translation of the mRNA component of CVnCoV in a rabbit reticulocyte lysate system. Translation of nascent proteins was detected via Western blotting. Water and luciferase control mRNA were employed as negative and positive controls, respectively. HeLa cells were transfected with 2 μg of the mRNA component of CVnCoV. 24h post-transfection, cells were analysed for S expression via (C) Western blotting using an S-specific antibody and (D) flow cytometric analyses using an S-specific antibody either with or without membrane permeabilisation allowing detection of total (intracellular) or cell surface bound (cell surface) S protein. Relative protein expression in Western blotting was quantified using the Image Studio Lite Ver 5.2 software. Geometric mean fluorescence intensity (GMFI) of transfected HeLa cells are expressed as mean + standard deviation (SD) of duplicate samples. All blots derive from the same experiment and that they were processed in parallel. NTD: N-terminal domain, RBD: receptor binding domain, IVT: *in vitro* translation, TM: transmembrane domain, Tub: Tubulin

To characterise mRNA-induced innate and adaptive immune responses, Balb/c mice were immunised with 2 μg of LNP-formulated mRNA encoding SARS-CoV-2 S-2P (CVnCoV). Innate responses were assessed in sera of CVnCoV vaccinated mice 14 hours post injection, the time point when mRNA-induced pro-inflammatory cytokines and chemokines peak^6^. These analyses demonstrated the induction of a balanced immune response upon CVnCoV injection that exhibited no bias towards IFN_γ_ or IL4, IL-5 and IL-13, indicative of a T_h_1 and T_h_2 response, respectively. Low levels of pro-inflammatory cytokines IL-6, IFNα were detectable in serum, while TNF and IL1β remained undetectable (Figure 2A).

**Figure 2:**
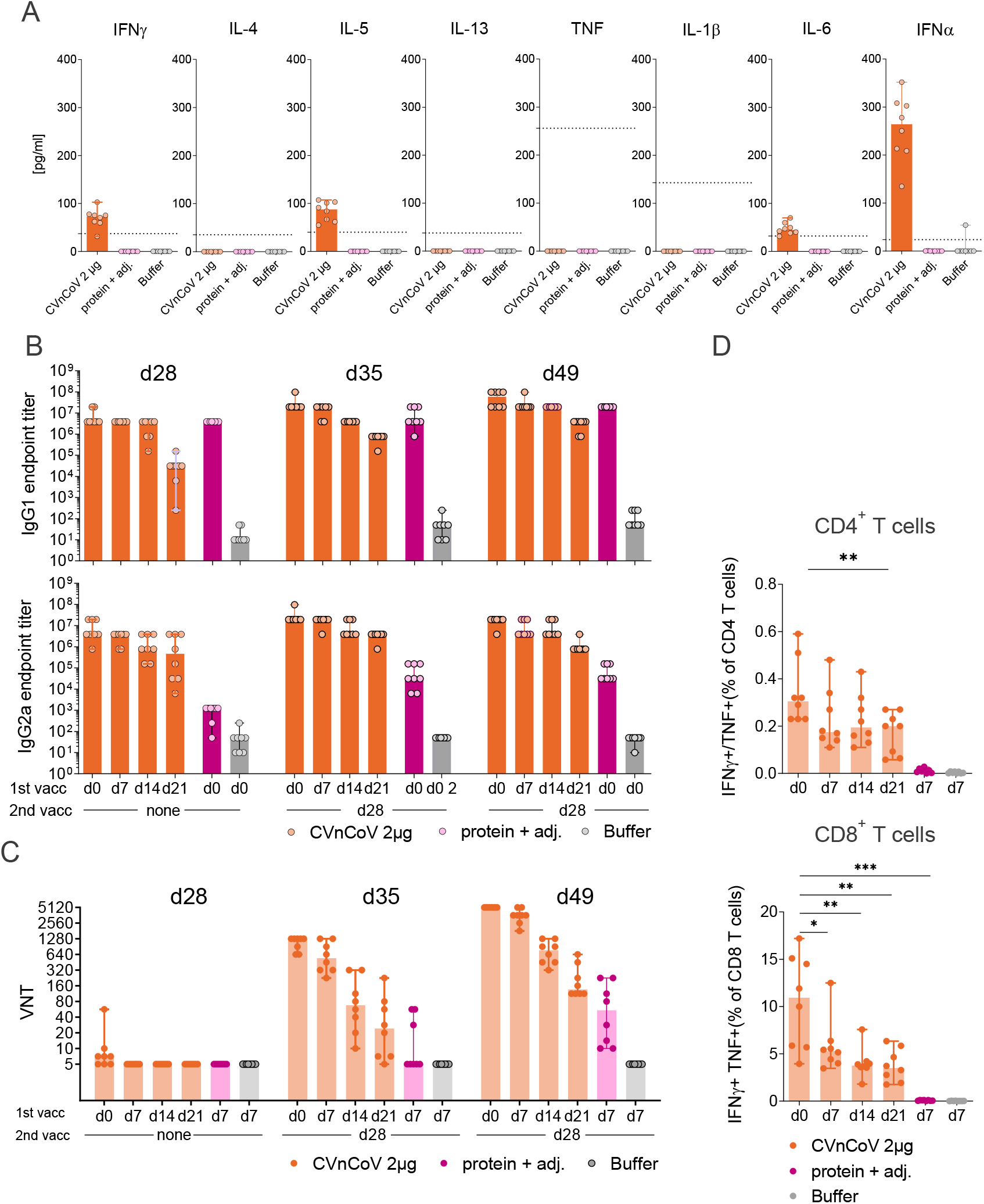
CVnCoV elicits high levels of humoral and cellular immune responses. Female Balb/c mice (n=8/group) were vaccinated IM on day 0, day 7, day 14 or day 21 with 2 μg of CVnCoV. All animals received a second immunization on day 28. 1.5μg of Alum adjuvanted SARS-CoV-2 S ectodomain (S_ECD_) protein and 0.9% NaCl (buffer) administered on day 0 and day 28 served as positive and negative control, respectively. (A) Cytokine induction in sera of vaccinated mice was assessed 14 h post injection. Dotted lines represent the lower limit of quantification (B) S_ECD_ protein specific binding antibodies, displayed as endpoint titres for IgG1 and IgG2a and (C) CPE-based virus neutralising titres in serum upon one (day 28) or two vaccinations (day 35 and day 49). (D) Multifunctional IFN-γ/TNF positive CD4^+^ T and CD8^+^ T cells analysed in splenocytes isolated on day 49. Cells were stimulated with a specific SARS-CoV-2 spike protein peptide library (15mers 11 overlapping) for 24h followed by intracellular cytokines staining and detection by flow cytometry. Each dot represents an individual animal, bars depict the median. Statistical analysis was performed using Mann-Whitney testing.

To determine immunization schedule impact, two injections of CVnCoV were given 4, 3, 2 or 1 week apart, i.e. on d0/d28, d7/d28, d14/d28 or d21/d28. Spike ectodomain (S_ECD_)-binding antibodies, found to correlate with the induction of virus neutralizing antibodies in COVID-19 patients^29^, developed rapidly upon a single injection with CVnCoV. Seven days after a single vaccination, robust IgG1 and IgG2a titres were present and increased over time (Figure 2B). Median endpoint titres on day 7 were 3.1 x 10^4^ and 4.7 x 10^5^, for IgG1 and IgG2a, respectively, and increased to 3.9 x 10^6^ for both IgG1 and IgG2a on d28. Second injection with CVnCoV induced increase in S_ECD_-binding IgG1 and IgG2a antibodies, while a trend towards lower immune responses induced by shorter intervals between first and second vaccination remained detectable until termination of the experiment in week 7 (d49). As expected, Alum-adjuvanted S_ECD_ protein domain, employed as a control for T_h_2 biased responses, induced high titres of IgG1 but comparatively low levels of IgG2a. Induction of virus neutralizing titres (VNTs) in mouse sera were assessed in a cytopathic effect (CPE)-based assay with wild-type SARS-CoV-2. Detectable levels of VNTs started to develop 4 weeks after a single injection, while a second injection led to significant VNT increase in all groups. Analyses one and three weeks after the second vaccination showed that responses rose over time (Figure 2C). By termination on day 49, all sera derived from animals vaccinated on d0/28 were able to neutralise 50% of infecting virus (100 TCID_50_) in cell culture at a dilution of 1:5120. For comparison, two serum samples from symptomatic convalescent subjects chosen for the presence of high levels of binding antibodies provided titres of 1:320 −1:640.

Strong induction of IFN_γ_/TNF double positive T cells in CVnCoV-vaccinated mice (Figure 2D) was demonstrated by T cell analyses via intracellular staining (ICS) of splenocytes isolated 3 weeks after the second vaccination (d49). T cell responses benefited from longer intervals between first and second vaccination. Highest responses were detected in animals vaccinated on d0/d28 with median values of IFN_γ_^+^/TNF^+^ CD4^+^ and CD8^+^ T cell of 0.34% and 10.5%, respectively. In contrast, IFN_γ_^+^/TNF^+^ CD4^+^ or CD8^+^ T cells remained undetectable in the protein control group.

Mice vaccinated with CVnCoV at 0.25 μg, 1 μg or 4 μg elicited dose dependent humoral responses. IgG1 and IgG2a antibodies induced by CVnCoV interacted with S_ECD_, the isolated S receptor binding domain (RBD), the trimeric form of the S-protein (S trimer), and with the isolated N-terminal domain (NTD) (Figure 3A). CVnCoV induced comparable levels of S_ECD_, RBD and trimeric S reactive antibodies, with the exception of the lowest 0.25μg dose, which generated lower RBD responses. Of note, antibodies directed against the NTD, an alternative target of VNTs next to the RBD^29^, were detectable for all doses but overall remained lower than levels of S_ECD_-, RBD- and S trimer-binding antibodies. As observed previously, VNTs started to develop 3 weeks after a single vaccination and robust levels were detectable after a second vaccination with titres of 1:2560 and 1: 5120 for the 1 μg and 4 μg dose groups, respectively. Even the lowest dose of 0.25μg induced detectable levels of VNTs of approximately 1:14 (Figure 3B). Further experiments demonstrated comparable relative avidity for S_ECD_ specific IgG1 and IgG2a antibodies (Figure 3C). CVnCoV-induced antibodies that efficiently competed with a neutralizing monoclonal antibody for binding to native SARS-CoV-2 S expressed on the cellular surface (Figure 3D).

**Figure 3:**
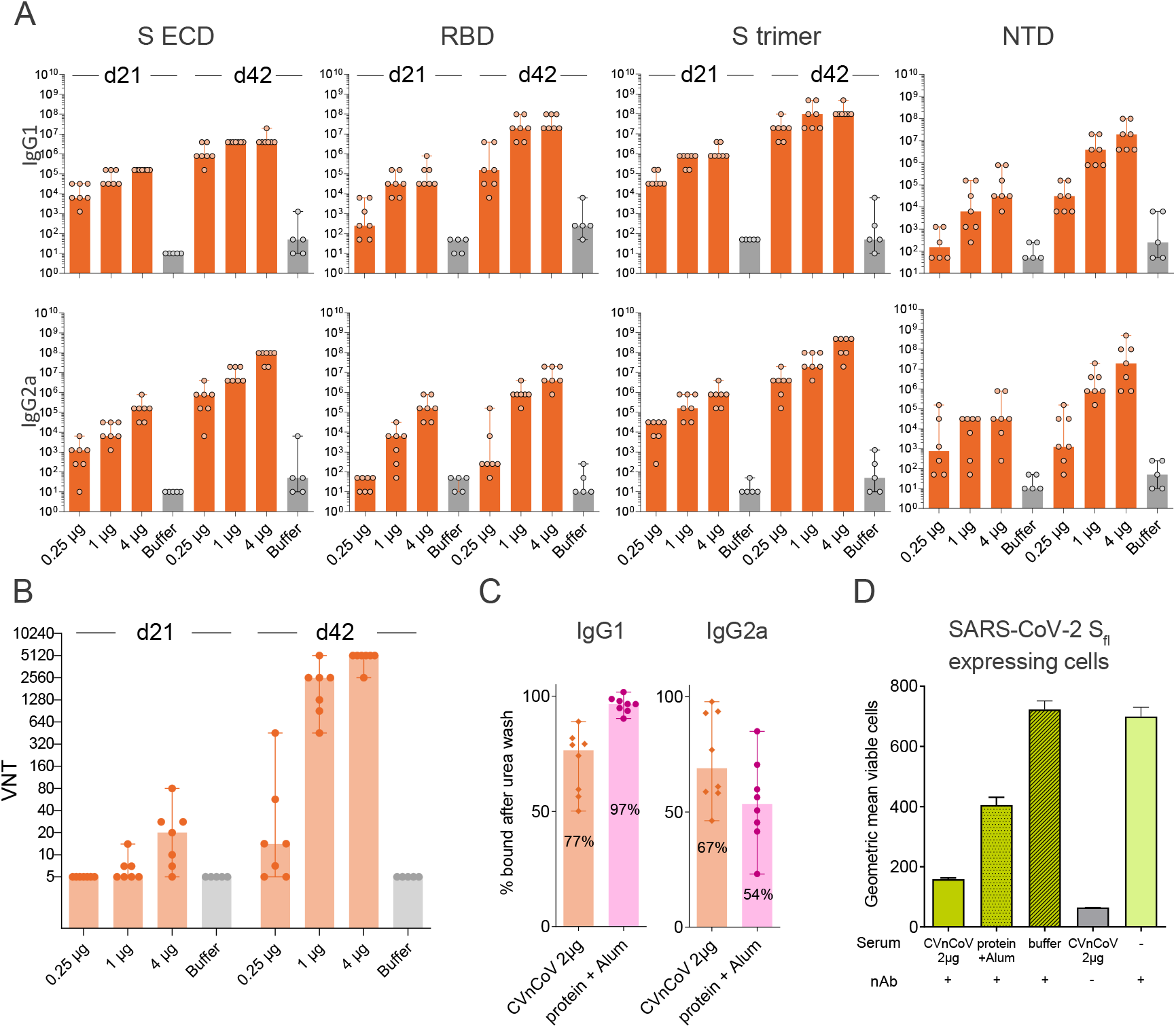
CVnCoV elicits high titres of functional antibodies against SARS-CoV-2. Female Balb/c mice (n=7/group) were vaccinated IM with 0.25, 1 and 4 μg of CVnCoV on day 0 and day 21. Animals (n=5/group) vaccinated with NaCl (Buffer) served as negative controls (A) Spike protein specific binding antibodies binding to S_ECD_, RBD, S trimer and NTD, displayed as ELISA endpoint titres for IgG1 and IgG2a for day 21 and day 42. (B) CPE-based VNTs in serum samples taken on day 21 and day 42. (C) Percent of S_ECD_ binding IgG1 and IgG2a antibodies remaining using serum from animals vaccinated with 2μg of CVnCoV or 1.5μg Alum adjuvanted S_ECD_ upon 30 min wash in 8% urea compared to buffer wash. (D) S-specific signal detectable on the surface of HeLa cells expressing full length SARS-CoV-2 S upon incubation with fluorescently labelled monoclonal neutralising antibody in the absence or presence of mouse serum. Serum employed was either from mice vaccinated with 2μg of CVnCoV, 1.5 μg of Alum adjuvanted protein or buffer. Each dot represents an individual animal, bars depict the median. S_ECD_: S ectodomain, RBD: receptor binding domain of S, NTD: N-terminal domain of S, nAb: neutralising antibody

Next, we sought to address the protective efficacy of CVnCoV in Syrian hamsters. Animals vaccinated with two CVnCoV doses of 2 μg or 10 μg in a 4-week interval developed dose-dependent S_ECD_ binding IgG antibodies after the first vaccination that increased upon the second (Figure 4A). Median endpoint titres of animals vaccinated with 10 μg of CVnCoV were 1.6 x 10^5^ after one dose and peaked at 7.8 x 10^5^ on day 42. Binding antibody titres were not assessed in animals previously infected with SARS-CoV-2. In agreement with mouse results generated in a similar assay with varied incubation time, CPE-based VNT analysis showed no significant levels after one vaccination in any of the CVnCoV vaccinated groups. However, two vaccinations with 10 μg of CVnCoV were able to induce robust levels of neutralising titres in hamsters. Of note, virus employed for this assay featured the D614G mutation, while CVnCoV encoded S does not include this mutation. The median titres peaked at 1:134 on day 42. For comparison, control animals infected with wild-type SARS-CoV-2 on day 0 developed VNTs of 1:191. A control group that received Alum-adjuvanted S_ECD_ protein developed IgG antibodies but undetectable VNT levels (Figure 4B).

**Figure 4:**
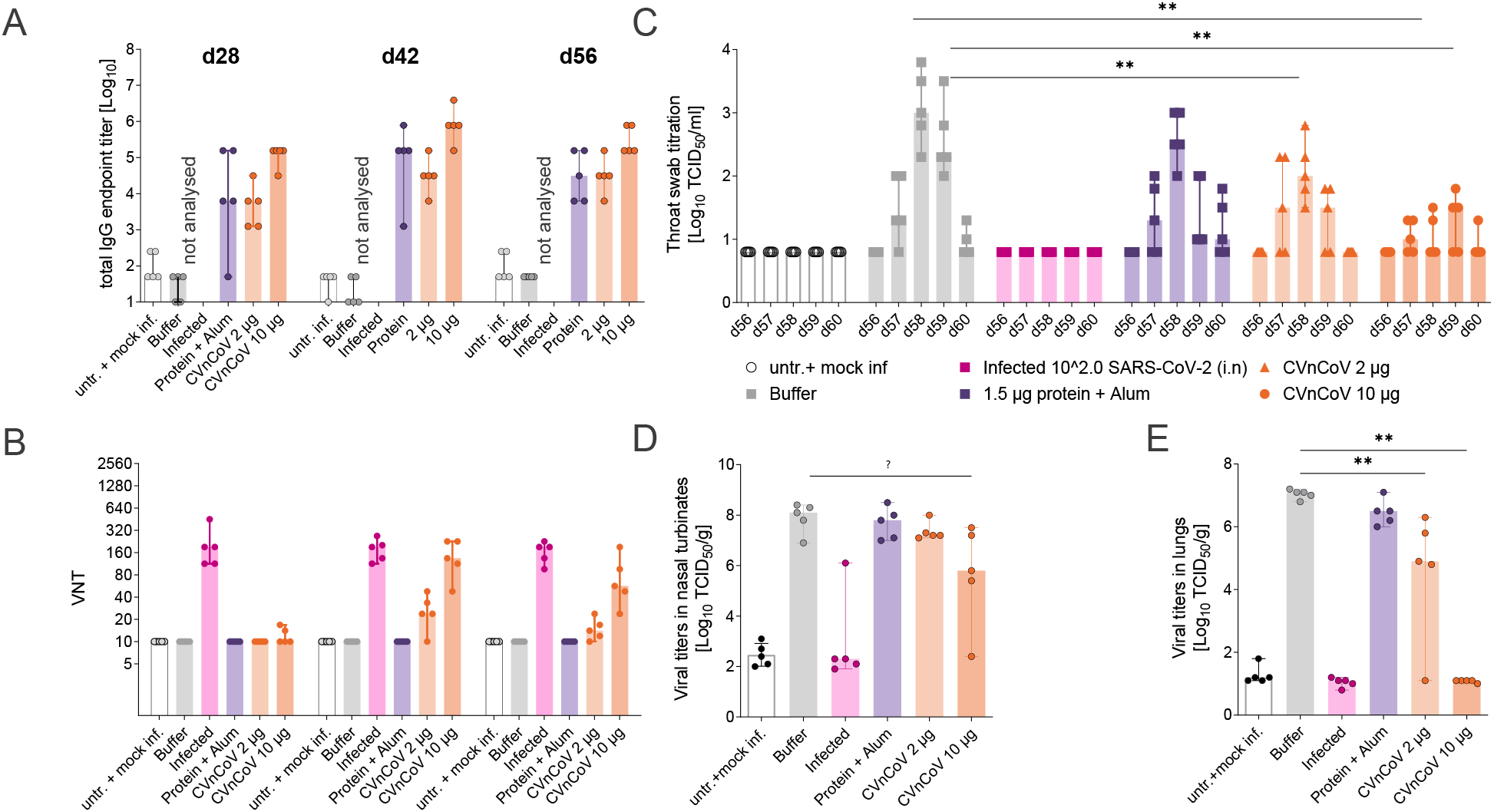
CVnCoV protects hamsters from SARS-CoV-2 challenge infection. Syrian golden hamsters (n=5/group) were vaccinated with 10 μg or 2 μg of CVnCoV, 1.5μg of Alum adjuvanted S_ECD_ protein or buffer on day 0 and d28. As additional controls, animals were either left untreated or infected intranasally (IN) with 10^2^ TCID_50_/dose of SARS CoV-2 on day 0 of the experiment. (A) Total IgG antibodies binding to S_ECD_ displayed as ELISA endpoint titres of all groups except for infected animals that were not analysed or (B) VNTs determined via CPE-based assay upon one (day 28) or two vaccinations (day 42 and day 56). On day 56, all animals except for the untreated group were challenged by IN infection with 10^2^ TCID_50_/dose of SARS CoV-2 in a total dose volume of 0.1 ml. Animals in the untreated groups were mock infected with buffer as a negative control. Animals were followed for four days post challenge (p.c.) and euthanised on day 60 of the experiment. Detectable levels of replication competent virus in (C) throat swabs on days 56 to day 60, (D) nasal turbinate on day 60 and (E) lung tissues on day 60 were analysed. Each dot represents an individual animal, bars depict the median. Statistical analysis was performed using Mann-Whitney testing.

On day 56, four weeks after the second vaccination, all animals were challenged with 10^2^ TCID_50_/dose of SARS-CoV-2 featuring D614G. While this infection dose is low compared to other studies that employed doses of 2 x 10^3^ to 2 x 10^6^ TCID_50_^30, 31, 32, 33, 34, 35, 36^, our results show robust levels of viral replication in the upper and lower respiratory tract (Figure 4) as well as typical weight loss for this model system^30, 31, 33^ (Supplementary Figure 1) in the buffer control group.

Analysis of animal weight upon challenge on days 56 to 60 revealed a maximal weight loss of 10% in buffer control animals with a steady decline throughout the observation period, while untreated and mock infected animals gained 3% of weight. All other groups, including animals previously infected with SARS-CoV-2, exhibited intermediate values of 5-6% on d60 compared to d56 with the weight drop stabilising on d59 (Supplementary Figure 1). Buffer control animal throat swabs, taken daily from day 56 to termination on day 60, showed peak replication-competent viral titres of approximately 10^3^ TCID_50_/ml two days post challenge. These titres were nearly undetectable on d60. Animals previously infected with SARS-CoV-2 remained negative throughout the experiment. Viral levels were significantly reduced in throat swabs of both CVnCoV-vaccinated groups. Vaccination with 10 μg of CVnCoV resulted in significantly diminished and delayed viral peaks at 10^1.5^ TCID_50_/ml three days post challenge. At least 2 out of 5 animals in this group remained negative throughout the testing period (Figure 4 C). Viral levels in nasal turbinates revealed less pronounced, but detectable dose-dependent reduction of viral replication (Figure 4D). Importantly, animals vaccinated with 10 μg of CVnCoV exhibited no detectable virus in the lungs, proving the ability of CVnCoV to protect animals from viral replication in the lower respiratory tract (Figure 4E).

Histopathological analyses demonstrated the occurrence of alveolar damage and moderate to severe inflammation of upper (nasal turbinates) and lower (alveoli, bronchi, trachea) respiratory tract in the buffer control group upon SARS-CoV-2 infection, proving the suitability of this model to assess virus-induced pathology. Consistent with protection from viral replication in the lungs, two vaccinations with 8 μg of CVnCoV significantly reduced histopathological changes in the lungs. Animals in this group featured significantly decreased inflammation of the lower respiratory tract, i.e. tracheitis, bronchitis, bronchiolitis and alveolitis. Importantly, a dose of 2 μg, which led to the induction of binding antibodies with low levels of VNTs, did not induce increased histopathology scores. Group comparisons for differential gene expression in lung homogenates showed no significant change in the induction of IL-4 or IL-5 in the mRNA groups compared to buffer or mock infection groups (data not shown). The presented data give some indication that Alum-adjuvanted protein vaccine, which induces a T_h_2 shifted immune response with no detectable levels of VNTs but high levels of binding antibodies, causes increased histopathology scores in hamsters (Figure 5).

**Figure 5:**
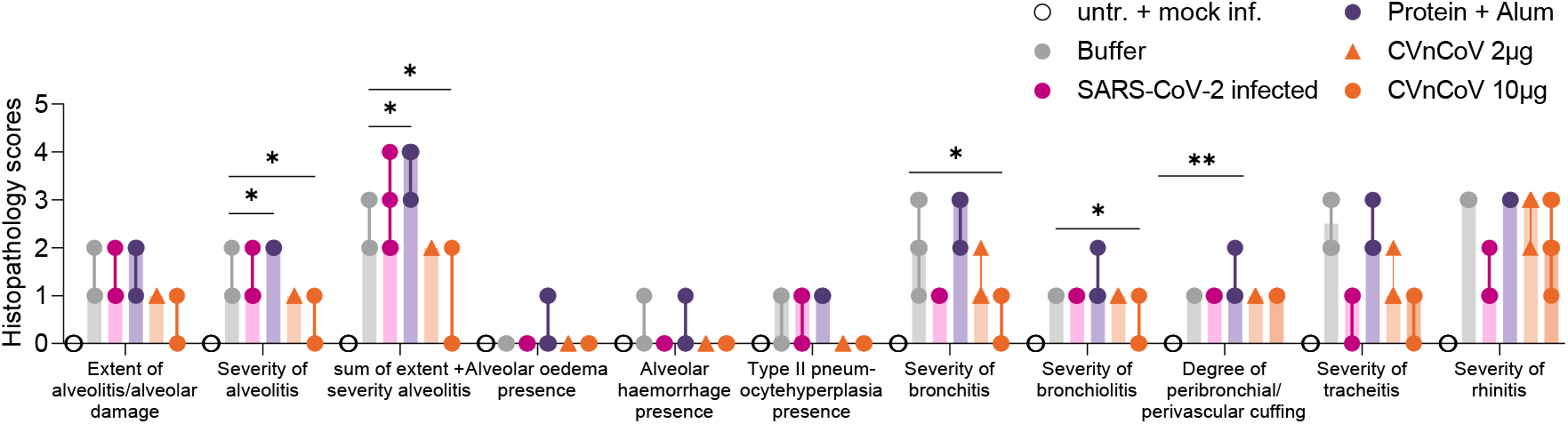
CVnCoV protects the respiratory tract from challenge infection with no signs of vaccine enhanced disease. Histopathological analyses of hamsters vaccinated with 10 μg or 2 μg of CVnCoV, 1.5μg of Alum adjuvanted S_ECD_ protein or buffer on day 0 and d28, left untreated and mock infected,or infected intranasally (IN) with 10^2^ TCID_50_/dose of SARS CoV-2 on day 0 of the experiment followed by of SARS CoV-2 challenge infection on d56. Histopathological analysis was performed on day 60, four days post challenge infection, on formalin-fixed, paraffin embedded tissues sections sampled on day 4 post challenge. Histopathological assessment scoring was performed according to severity of inspected parameter. Each dot represents an individual animal, bars depict the median, Statistical analysis was performed using Mann-Whitney testing.Supplementary Materials

In order to exclude that potential pathological changes upon CVnCoV vaccination were induced but remained undetected at day 4 post challenge, a parallel experiment in hamsters terminated on day 7 was conducted. This later time point prevented the determination of viral levels in lungs which were undetectable in all groups (data not shown) but provided overall higher levels of pathological changes in the respiratory tract. The analysis of median gross pathology revealed values of 30% affected lung tissue in buffer control animals. In contrast, all and 3/5 animals in the 8 μg and 2 μg CVnCoV vaccinated groups, respectively, featured no visually detectable levels of pathological changes of the lungs (Supplementary Figure 2A). In agreement with analyses on day 4 post challenge, histopathology performed on day 7 post challenge demonstrated significant reduction of inflammation in both of upper and lower respiratory tract samples in both CVnCoV vaccinated groups with no indication of increased inflammation in the lower dose group (Supplementary Figure 2B).

## Discussion

Here we show that CVnCoV mRNA is translated into a protein that was cleaved, post-translationally modified, and presented on the cell surface, mimicking viral protein presentation during infection. In line with previous results^6^, CVnCoV induced innate immune responses, including IL-6 and IFNα in mice, indicative of a pro-inflammatory environment for the induction of cellular and humoral immune responses. Incorporation of two prolines that prevent structural rearrangement to the post-fusion state of coronavirus S proteins^25,24^ in the CVnCoV design was aimed to increase S-induced immune responses as observed in the context of MERS^24^. Indeed, vaccination with CVnCoV induced robust humoral responses featuring high levels of virus neutralizing titers in mice and hamsters that were dose- and vaccination schedule-dependent. Antibody responses in mice developed rapidly and to comparatively high levels^37^ upon injection and were reactive with both S domains targeted by human VNTs, i.e. the RBD domain as the primary target of neutralizing antibodies^38,39^, as well as the NTD^40^. CPE-based detection of virus neutralizing antibody responses revealed that CVnCoV induced high levels of VNTs upon two injections against both viruses featuring D and G on position 614 in S for mice and hamsters, respectively. In hamsters, VNTs were comparable to titres induced upon natural infection with wild type SARS-CoV-2. As previously shown for mRNA vaccines against SARS-CoV-2^13,37^, CVnCoV was able to induce both S-specific CD4^+^ and CD8^+^ T cell responses in mice that were comparatively high at low vaccine doses^37^. T cells are known to contribute to protection in respiratory infections, with virus-specific CD8^+^ T cells representing effectors of viral clearance from lung tissue^41^. In a clinical study, influenza-specific CD8^+^ T cells protected against symptomatic influenza infection in the absence of protective antibodies^42^. Furthermore, memory CD8^+^ T cells provided substantial protection in a lethal SARS-CoV-1 mouse model^43^. In the context of SARS-CoV-2, recent data have demonstrated that CD8+ T cells contribute to viral control in a rhesus macaque model. Data presented here demonstrate that CVnCoV vaccination induces high levels of antigen-specific CD8^+^ IFN_γ_/TNF double-positive T cells likely to contribute to vaccine efficacy upon viral challenge.

Consistent with robust immune responses, CVnCoV protected hamsters against SARS-CoV-2 viral challenge featuring the D614G mutation in S, proving CVnCoV’s ability to protect against the most prevalent form of the virus. These experiments showed significant reduction in replicating virus levels in the upper respiratory tract and the absence of detectable live virus in the lungs of animals upon two vaccinations with 10 μg of CVnCoV. This result is in line with previous observations in other respiratory viruses as well in SARS-CoV-2 challenge models in hamsters^30^ and non-human primates^13,44^, where robust immune responses induced better protection in the lower than in the upper respiratory tract.

Importantly, data presented here support a favourable safety profile of CVnCoV. A potential risk of vaccine development is the induction of vaccine-enhanced disease or pathology, either induced by antibodies (antibody-dependent enhanced disease, ADE) or vaccine-associated enhanced respiratory disease (VAERD) (reviewed in^45,46,45,47^)· ADE, that leads to exacerbated disease by enhancement of viral entry, has been described for a feline coronavirus^48^ as well as for other viruses such as respiratory syncytial virus (RSV) and measles virus. ADE has largely been associated with the presence of nonneutralizing antibodies (reviewed in^47^). VAERD that occurred in the context of whole-inactivated vaccines^49, 50^ has been linked to inflammation induced by T_h_2-biased immune responses and high ratios on non-neutralising to neutralising antibodies due to presentation of conformationally incorrect antigens (reviewed in^46^). Vaccine-enhanced disease has been described in non-clinical models for SARS and MERS-CoV (reviewed in^51,52^). There is no evidence of vaccine-enhanced disease in animal models of SARS-CoV-2^44, 12, 53, 54^, nor clinical evidence that this might be a concern in SARS-CoV-2 infections. However, such hypothetical effects need to be carefully excluded during vaccine development. Several lines of evidence suggest that the risk of CVnCoV to induce vaccine-enhanced disease is low. Firstly, immunogenicity data generated in two animal models demonstrated robust levels of VNTs compared to binding antibodies upon vaccination with CVnCoV. Secondly, the vaccine elicited a balanced T_h_1/T_h_2 profile, indicated by the induction of equivalent levels of IgG1 and IgG2a antibodies which exhibited comparable relative avidity to SARS-CoV-2 S as well as a cytokine profile in mice that gave no indication of a T_h_2 bias, i.e. induction of IL4, IL5 and IL13. In addition, challenge infection in hamsters did not reveal evidence of disease enhancement, neither by enhanced viral levels in the respiratory tract, nor inflammation of lung tissue, which was assessed at both early (d4) and late (d7) time points post challenge. This observation remained true in a group of animals vaccinated with 2μg, a suboptimal dose of CVnCoV that induced high levels of binding antibodies in the absence of significant virus neutralizing titres. Here, viral challenge led to breakthrough viral replication in the lungs. However, neither viral levels nor tissue inflammation on d4 or d7 was enhanced giving strong support for a favourable safety profile of CVnCoV.

We conclude that CVnCoV represents a promising vaccine candidate against SARS-CoV-2 in terms of potency and safety. The presented data supported the start of CVnCoV clinical trials. Phase 1 and 2 evaluation of CVnCOV is currently ongoing and data are collected to initiate pivotal phase 3 clinical testing.

## Material and Methods

### mRNA vaccines

The mRNA vaccine is based on the RNActive^®^ platform (claimed and described in e.g. WO2002098443 and WO2012019780)) and is comprised of a 5’ cap structure, a GC-enriched open reading frame (ORF), 3’ UTR, polyA tail and does not include chemically modified nucleosides. Lipid nanoparticle (LNP)-encapsulation of mRNA was performed by Acuitas Therapeutics (Vancouver, Canada) or Polymun Scientific Immunbiologische Forschung GmbH (Klosterneuburg, Austria). The LNPs used in this study are particles of ionizable amino lipid, phospholipid, cholesterol and a PEGylated lipid. The mRNA encoded protein is based on the spike glycoprotein of SARS-CoV-2 NCBI Reference Sequence NC_045512.2, GenBank accession number YP_009724390.1 and encodes for full length S featuring K_986_P and V_987_P mutations.

### *In vitro* protein expression

*In vitro* translation was performed using Rabbit Reticulocyte Lysate System, Nuclease Treated (Promega, Cat. L4960) according to the manufacturer’s instructions. In brief, 2 μg of mRNA encoding for SARS-CoV-2 S or control mRNA encoding for luciferase were denatured for 3 minutes at 65°C and incubated with Rabbit Reticulate Lysate, Amino Acid Mixtures, Transcend™ Biotin-Lysyl-tRNA (Promega, Cat.L5061), RNasin^®^ Ribonuclease Inhibitors (Promega, Cat. N2511) and Nuclease-Free water in a total volume of 50μl at 30°C for 90 minutes. *In vitro* translated proteins in 1x Lämmli buffer were proteins were separated on 4-20% Mini-PROTEAN^®^ TGX™ Precast Protein Gels (BioRad) and transferred to an Immobilon-FL PVDF membrane (Merck, Millipore, Cat. IPFL00010). Biotinylated Lysyl residues incorporated in the nascent proteins were detected using IRDye^^®^^ 800CW Streptavidin (LICOR, Cat. 926-32230). Protein detection and image processing were carried out in an Odyssey^^®^^ CLx Imaging system and LI-COR’s Image Studio version 5.2.5 according to manufacturer’s instructions.

For detection of mRNA expression in cell culture, HeLa cells were seeded in 6-well plates at a density of 400.000 cells/well. 24h later, cells were transfected with 2μg of mRNA per well via Lipofection. For this, RNAs were complexed with Lipofectamine 2000 (Life Technologies) at a ratio of 1:1.5 and transfected into cells according to manufacturer’s protocol. Protein expression was assessed 24h post transfection. For Western blotting, cells were lysed in 1x Lämmli buffer, proteins were separated on 4-20% Mini-PROTEAN^®^ TGX™ Precast Protein Gels (BioRad) and transferred to a nitrocellulose membrane (Odyssey nitrocellulose membrane 0.22 μm (Li-Cor, Cat 926-31092). Specific proteins were detected using rabbit anti-SARS Spike c-terminal (Abcam, Cat. ab252690) and mouse anti tubulin (Abcam, Cat. ab7291), followed by goat anti-rabbit IgG IRDye^®^ 800CW (Li-Cor, Cat. 926-32211) and goat anti-mouse IgG IRDye^®^ 680RD (Li-Cor, Cat. 926-68070), respectively. Protein detection and image processing were carried out in an Odyssey^®^ CLx Imaging system and LI-COR’s Image Studio version 5.2.5 according to manufacturer’s instructions.

For FACS analysis, cells were fixed and analysed with intact (surface staining) or permeabilised plasma membranes via treatment with Perm/Wash buffer (BD Biosciences, Cat. 554723) (intracellular staining). Specific S protein expression was assessed via staining with Human anti SARS CoV S antibody (CR3022) (Creative Biolabs, Cat. MRO-1214LC) followed by goat anti-human IgG F(ab’)2 fragment PE antibody (Immuno Research, Cat. 109-116-097) in a BD FACS Canto II cell analyzer and the FlowJo 10 software.

### Animals

Mice (BALB/c, 8-12 weeks of age) were obtained from Janvier Laboratories (Le Genest-Saint-Isle, France). For internal studies, experiments were approved by the Regional council Tübingen, animal testing license CUR 03/20. For studies performed externally, Balb/c mice were provided and handled by Preclinics Gesellschaft für präklinische Forschung mbH, (Potsdam, Germany). All animal experiments were conducted in accordance with German laws and guidelines for animal protection and appropriate local and national approvals.

Syrian golden hamsters, 11 to 13 weeks old, were obtained from Envigo (Indianapolis, IN, United States). Animals were housed and all procedures were performed by Viroclinics Xplore animal facility (Schaijk, The Netherlands) under conditions that meet the standard of Dutch law for animal experimentation and in accordance with Directive 2010/63/EU of the European Parliament and of the Council of 22 September 2010 on the protection of animals used for scientific purposes. Ethical approval for the experiment was registered under protocol number AVD277002015283-WP08.

### Immunisations

Animals were injected intramuscularly (IM) with LNP-formulated SARS-CoV-2 mRNAs (CVnCoV), NaCl as negative control, 1μg of formalin and heat inactivated SARS-CoV 2 adjuvanted in Alhydrogel (Brenntag) 1% or 1.5 μg of recombinant SARS-CoV-2 spike protein (S1+S2 ECD, His tag; Sino Biological, Cat. 40589-V08B1) adjuvanted in Alhydrogel (Brenntag) 2% as positive controls,. As an additional positive control, hamsters were infected intranasally (IN) with 10^2^ TCID_50_/dose of SARS CoV-2 isolate BetaCoV/Munich/BavPat1/2020 in 0.1 ml on d0 of the experiment as indicated.

### Challenge infections

Animals were challenged by infection IN with 10^2^ TCID_50_/dose of SARS CoV-2 (BetaCoV/Munich/BavPat1/2020) in a total dose volume of 0.1 ml on day 56, four weeks following the second immunisation. Animals were followed for four or seven days post challenge (p.c.) and euthanised on day 60 or day 63 of the experiment as indicated.

### Antibody analysis

Blood samples were taken via retro-orbital bleeding and SARS-CoV-2 S-specific IgG1 and IgG2a antibodies (mice) or total IgG (hamster) antibodies were detected via ELISA. Plates were coated with 1μg/ml of SARS-CoV-2 spike protein (S1+S2 ECD, His tag; Sino Biological, Cat. 40589-V08B1), SARS-CoV-2_DB_his (monomeric RBD) (Institute for Protein Design, DT# 35961, provided via GH-VAP), trimeric S protein, i.e. SARS-CoV-2_Secto_FO-His (Spike) (Institute for Protein Design, DT# 35962, provided via GH-VAP) or recombination SARS-CoV-2 spike N-terminal domain protein (NTDv1-Hisb3) (provided by Ian Wilson, The Scripps Research Institute via GH-VAP) for 4-5h at 37°C. Plates were blocked overnight in 1% BSA (mice) or 10% milk (hamsters), washed and incubated with serum for 2h at room temperature. For detection, mouse sera were incubated with biotin rat anti-mouse IgG1 (BD Pharmingen, Cat. 550331) or biotin rat anti-mouse IgG2a (BD Pharmingen, Cat. 550332), hamster sera with biotin goat anti-hamster (Syrian) IgG antibody (BioLegend, Cat: 405601) followed by incubation with HRP-Streptavidin (BD, Cat: 554066). Detection of specific signals was performed in a BioTek SynergyHTX plate reader, with excitation 530/25, emission detection 590/35 and a sensitivity of 45.

To assess binding competition of sera from vaccinated animals with neutralizing mAb, HeLa cells were transfected with mRNA encoding for SARS-CoV-2 S protein as described above. 24 h post transfection, approximately 160,000 cells/well were transferred into a 96-well V-bottom plate and incubated with 100 μl of 1:10 diluted mouse serum from mice vaccinated with CVnCoV, alum adjuvanted protein or buffer, at 4°C overnight. As a positive control, an additional sample of transfected cells remained untreated with serum. The cells were then incubated with 200 ng/well of R-PE labelled rabbit anti-SARS-CoV-2 spike neutralizing monoclonal antibody (Sino Biological, Cat: 40592-R001) at 4°C for 2 h. As a negative control, a sample of transfected cells treated with serum from CVnCoV vaccinated animals was incubated with buffer without labelled antibody. Antibody labelling with R-PE using the Zenon™ labelling kit for rabbit IgG (Invitrogen, Cat: Z25355) was performed according to the manufacturer’s instructions. Specific binding of R-PE labelled monoclonal antibody to S protein expressing HeLa cells was assessed via FACS analysis as described above.

For the analysis of virus neutralizing titres of mouse sera, serial dilutions of heat-inactivated sera (56°C for 30 min) tested in duplicates with a starting dilution of 1:10 followed by 1:2 serial dilutions were incubated with 100 TCID_50_ of wild type SARS-CoV-2 (strain 2019-nCov/Italy-INMI1) for 1 hour at 37°C. Every plate contained a dedicated row (8 wells) for cell control which contains only cells and medium, and a dedicated row of virus control which contain only cells and virus. Infectious virus was quantified upon incubation of 100 μl of virus-serum mixture with a confluent layer of Vero E6 cells (ATCC, Cat.1586) followed by incubation for 3 days at 37°C and microscopical scoring for CPE formation. A back titration was performed for each run in order to verify the correct range of TCID_50_ of the working virus solution. VN titres were calculated according to the method described by Reed & Muench. If no neutralization was observed (MNt <10), an arbitrary value of 5 was reported. Analyses were carried out at VisMederi srl (Siena, Italy).

VN antibody titres of hamster serum samples were analysed upon heat inactivation of samples for 30 min at 56°C. Triplicate, serial two-fold dilutions were incubated with 10^2^ TCID_50_/well SARS-CoV-2 virus for one hour at 37°C leading to a sample starting dilution of 1:10. The virus-serum mixtures were transferred to 96 well plates with Vero E6 cell culture monolayers and incubated for five days at 37°C. Plates were then scored using the vitality marker WST8 and (100% endpoint) VN titres were calculated according to the method described by Reed & Muench. Analyses were was done at Viroclinics Xplore (Schaijk, The Netherlands).

### T cell analysis

The induction of antigen-specific T cells was determined using intracellular cytokine staining (ICS). For ICS, splenocytes from vaccinated and control mice were isolated and single cell suspensions were prepared in supplemented medium. 2×10^6^ splenocytes (200 μl) per well were stimulated for 5-6 h at 37 °C using an SARS-CoV-2 peptide library (JPT, PM-WCPV-S2) at 0.5 μg/ml. After 1 h Golgi Plug (BD Biosciences, Cat: 555029) was added in a dilution of 1:200 (50 μl) to the splenocytes to inhibit the secretion of intracellular cytokines. After stimulation, splenocytes were centrifuged, resuspended in supplemented medium and stored at 4 °C overnight. Following this, splenocytes were washed twice in PBS and stained with AquaDye (Invitrogen, Cat: L34957) solution at 4 °C for 30 min. After an additional washing step in FACS buffer (PBS with 0.5% BSA) cells were surface stained for Thy1.2, CD4 and CD8 and incubated with FcyR-block for 30 min at 4 °C in FACS buffer. After surface staining, splenocytes were washed in FACS buffer and fixed using Cytofix/Cytoperm (BD Biosciences) according to the manufacturer’s instructions. After fixation, splenocytes were washed in perm buffer and stained for IFN-/ and TNF for 30 min at 4 °C. After staining, the cells were washed with perm buffer, resuspended in FACS buffer supplemented with 2mM EDTA and 0.01% Natriumacid and stored at 4 °C until analysis. Splenocytes were analysed on a Canto II flow cytometer (BD Biosciences). Flow cytometry data were analyzed using FlowJo software (Tree Star, Inc, Ashland, USA.).

The following antibodies were used for flow cytometry analysis: anti-Thy1.2 FITC (clone 53-2.1; Biolegend, Cat.14304), anti-CD4 V450 (clone RM4-5; BD Biosciences, Cat. 560468), anti-CD8a APC-H7 (clone 53-6.7; BD Biosciences, Cat. 560182), anti-IFNy APC (clone XMG1.2, BD Biosciences, Cat. 554413) and anti-TNF PE (clone MP6-XT22, eBioscience, Cat. 25-7321-82).

### Cytokine analysis

Blood samples were taken via retro-orbital bleeding 14h after administration of CVnCoV or buffer. Serum cytokines (IFN-γ, IL-1β, TNF, IL-6, IL-4, IL-5 and IL-13) were assessed using cytometric bead array (CBA) using the BD FACS CANTO II. Serum was diluted 1:4 and BD Bioscience mouse cytokine flex sets were used according to manufacturer’s protocol to determine serum cytokine levels.

The following flex sets were used: Mouse IFN-γ Flex Set RUO (A4) (BD Bioscience, Cat. 558296); Mouse Il-13 Flex Set RUO (B8) (BD Bioscience, Cat. 558349); Mouse IL-1β Flex Set RUO (E5) (BD Bioscience, Cat. 560232); Mouse Il-4 Flex Set RUO (A7) (BD Bioscience, Cat. 558298); Mouse Il-5 Flex Set RUO (A6) (BD Bioscience, Cat. 558302); Mouse IL-6 Flex Set RUO (B4) (BD Bioscience, Cat. 558301); Mouse TNF Flex Set RUO (C8) (BD Bioscience, Cat. 558299).

IFN-α was quantified using VeriKine-HS Mouse IFNα Serum ELISA Kit (pbl, Cat. 42115-1) according to manufacturer’s instructions. Sera were diluted 1:100 and 50μl of the dilution was tested.

### Viral load in the respiratory tract

Detectable levels of replication competent virus in throat swabs, lung and nasal turbinate tissues post challenge were analysed. Quadruplicate, 10-fold serial dilutions were transferred to 96 well plates with Vero E6 cell culture monolayers and incubated for one hour at 37°C. Cell monolayers were washed prior to incubation for five days at 37°C. Plates were then scored using the vitality marker WST8 and viral titers (Log10 TCID_50_/ml or /g) were calculated using the method of Spearman-Karber. Analyses were done at Viroclinics Xplore (Schaijk, The Netherlands).

### Histopathology upon challenge in hamsters

Histopathological analysis was performed on tissues sampled on day 4 post challenge. After fixation with 10% formalin, sections were embedded in paraffin and the tissue sections were stained with haematoxylin and eosin for histological examination. Histopathological assessment scoring is as follows: Alveolitis severity, bronchitis/bronchiolitis severity: 0 = no inflammatory cells, 1 = few inflammatory cells, 2 = moderate number of inflammatory cells, 3 = many inflammatory cells. Alveolitis extent, 0 = 0%, 1 = < 25%, 2 = 25-50%, 3 = >50%. Alveolar oedema presence, alveolar haemorrhage presence, type II pneumocyte hyperplasia presence, 0 = no, 1 = yes. Extent of peribronchial/perivascular cuffing, 0 = none, 1 = 1-2 cells thick, 2 = 3-10 cells thick, 3 = >10 cells thick. Histopathological analysis was performed by Judith van den Brand DVM PhD dipl ECVP (Division of Pathology, Faculty of Veterinary Medicine, Utrecht University, The Netherlands).

### Gross lung pathology

At the time of necropsy, gross pathology was performed on each animal and all abnormalities were described. All lung lobes were inspected and the percentage of affected lung tissue evaluated via visual inspection and scoring determined.

## Data availability statement

The authors declare that all relevant data supporting the findings of this study are available within the paper and its supplementary information files.

## Acknowledgements

Thanks to Amy Shurtleff, Arun Kumar and Gerald Voss (CEPI) for their help and support setting up and evaluating experiments and for critically reading the manuscript.

Thanks to Jennifer Oduro and Jonas Fuener (preclinics mbH) for their support with mouse immunogenicity studies, their outstanding work, commitment and expertise.

Thanks to Koert Stittelaar and Kate Guilfoyle (Viroclinics) for performing the hamster challenge experiments and for their expert advice and excellent work throughout the project.

Special thanks to Prof. Emanuele Montomoli, Giulia Lapini and the Vismederi team for performing excellent work on virus neutralizing antibody assays.

Thanks to Acuitas Therapeutics and Polymun Scientific for LNP formulations.

Very special thanks to Julia Jürgens and Julia Schröder for their outstanding work performing mouse experiments and binding antibody analyses, Annette Moebes, Jessica Lahm, Michaela Trapp, Nina Schneck and Rebecca Winter for their relentless support in the lab. Andreas Theß, Moritz Thran, Wolfgang Große for their support with generating mRNA constructs, Stefanie Sewing, Susanne Braeuer and Aniela Wochner for mRNA production, Michael Firgens and Cibele Gaido for all their excellent work on project planning and communication and thanks to Igor Splawski for critically reading the manuscript.

## Competing interests

M.F-M. is a management board member and employee of CureVac AG, Tuebingen Germany. S.R., B.P., N.R., K.S. and S.O.M. are employees of CureVac AG, Tuebingen Germany, a publically listed company developing RNA-based vaccines and immunotherapeutics. All authors may hold shares or stock options in the company. S.R., B.P., N.R., K.S., M.F-M. inventors on several patents on mRNA vaccination and use thereof.

## Author contributions

S.R., M. M-F., S.O.M. and B.P. conceived and conceptualised the work and strategy. S.R. and B.P. designed *in vitro* and *in vivo* studies including hamster challenge studies. S.R., K.S. and N.A. analysed and interpreted data. S.R. wrote the manuscript. All authors supported the review of the manuscript.

## Funding

Work submitted in this paper was funded by the Coalition for Epidemic Preparedness Innovations (CEPI)

---

**Supplementary Figure 1:**
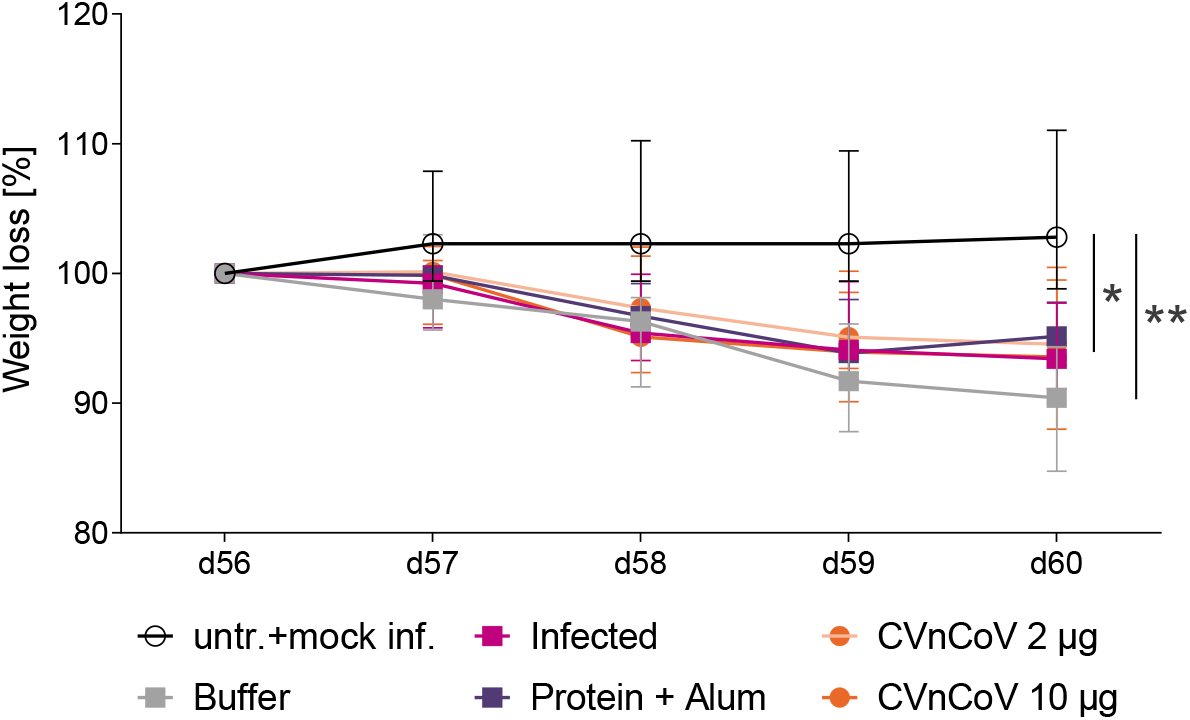
Weight loss upon challenge infection. Development of weight upon challenge infection on d56 was assessed in all groups, i.e. animals vaccinated with 10 μg or 2 μg of CVnCoV, 1.5μg of Alum adjuvanted S_ECD_ protein or buffer on day 0 and d28, left untreated and mock infected,or infected intranasally (IN) with 10^2^ TCID_50_/dose of SARS CoV-2 on day 0 of the experiment. Values are shown as median with range in % of value on d56 for all groups, statistical analysis was performed using Mann-Whitney testing.

**Supplementary Figure 2:**
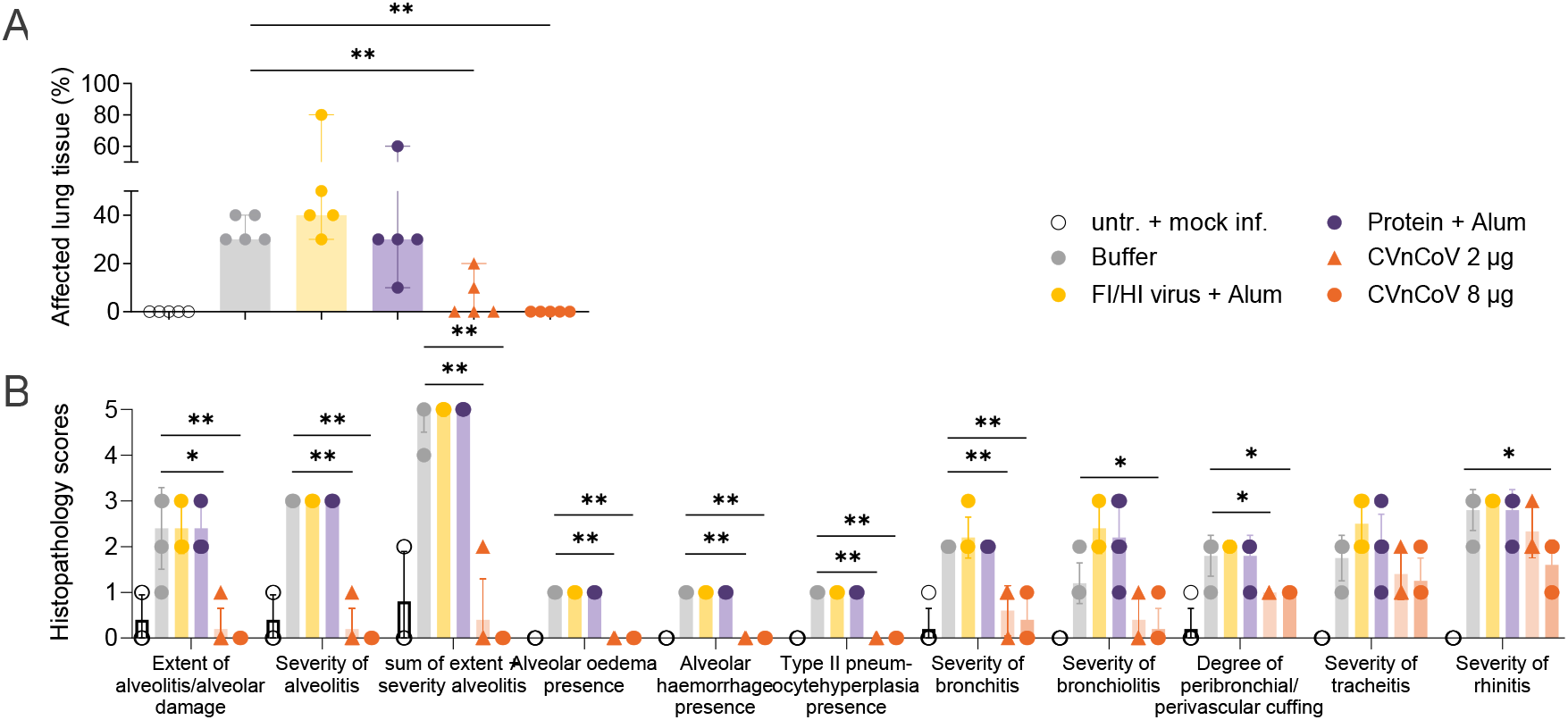
Vaccination with CVnCoV protects hamsters from pathological changes of the respiratory tract on day 7 post challenge. (A) Hamsters vaccinated with 8 μg or 2 μg of CVnCoV, 1μg of Alum adjuvanted, formalin and heat inactivated SARS-CoV-2, 1.5μg of Alum adjuvanted S_ECD_ protein or buffer on day 0 and d28, followed by of SARS CoV-2 challenge infection on d56 or left untreated and mock infected were assessed for gross pathological changes of the lungs on day 7 post challenge. Changes were determined via complete macroscopic post-mortem examination of all lung lobes. The percentage of affected lung tissue determined via visual inspection and scoring is shown. (B) Histopathological analysis was performed on day 63, seven days post challenge infection, on formalin-fixed, paraffin embedded tissues sections sampled on day 7 post challenge. Histopathological assessment scoring was performed according to severity of inspected parameter. Each dot represents an individual animal, bars depict the median, statistical analysis was performed using Mann-Whitney testing.

## Notes

### Summary of Updates

Additional data on CVnCoV protective efficacy in hamsters

